# Vein-to-blade ratio is an allometric indicator of climate-induced changes in grapevine leaf size and shape

**DOI:** 10.1101/2020.05.20.106906

**Authors:** Daniel H. Chitwood, Joey Mullins, Zoë Migicovsky, Margaret Frank, Robert VanBuren, Jason P. Londo

**Affiliations:** Department of Horticulture, Michigan State University, East Lansing, Michigan, 48824, USA; Department of Computational Mathematics, Science & Engineering, Michigan State University, East Lansing, Michigan, 48824, USA; Department of Plant, Food and Environmental Sciences, Faculty of Agriculture, Dalhousie University, Truro, NS B2N 5E3, Canada; School of Integrative Plant Science, Plant Biology Section, Cornell University, Ithaca, New York, 14850, USA; U.S. Department of Agriculture, Agriculture Research Service, Grape Genetics Research Unit, Geneva, New York 14456, US

**Keywords:** allometry, ampelography, ampelometry, grapevine, leaf morphology, leaf shape, leaf size, Vitaceae, Vitis, climate

## Abstract

**Premise:** As a leaf expands, its shape dynamically changes. Previously, we documented an allometric relationship between vein and blade area in grapevine leaves. Larger leaves have a smaller ratio of primary and secondary vein area relative to blade area compared to smaller leaves. We sought to use allometry as an indicator of leaf size to measure the environmental effects of climate on grapevine leaf morphology.

**Methods:** We measure the ratio of vein-to-blade area in 8,412 leaves from the same 208 vines across four growing seasons (2013, 2015, 2016, and 2017) using 21 homologous landmarks. Matching leaves by vine and node, we correlate size and shape of grapevine leaves with climate variables.

**Key results:** Vein-to-blade ratio varies strongly between years in ways that blade or vein area do not. Maximum daily temperature and to a lesser degree precipitation are the most strongly correlated climate variables with vein-to-blade ratio, indicating that smaller leaves are associated with heat waves and drought. Leaf count and overall leaf area of shoots and the vineyard population studied also diminish with heat and drought. Grapevine leaf primordia initiate in buds the year prior to when they emerge, and we find that climate during the previous growing season exerts the largest statistical effects over these relationships.

**Conclusions:** Our results demonstrate the profound effects of heat and drought on the vegetative morphology of grapevines and show that vein-to-blade ratio is a strong allometric indicator of the effects of climate on grapevine leaf morphology.

## INTRODUCTION

The morphology of grapevine (*Vitis* spp.) leaves uniquely allows their shape to be measured using geometric methods. Nearly all leaves in the genus *Vitis*, whether simple or compound, possess seven major veins with established nomenclature (OIV, 2018): a midvein (or L1), two distal veins (or superior, L2), two proximal veins (or inferior, L3), and two petiolar veins (or L4) that branch off the proximal veins forming a petiolar sinus encircling the petiolar junction. These veins also define the distal (or superior) and proximal (or inferior) sinuses of the leaf (Fig. 1). Ampelometry (“vine” + “the process of measuring”) has long recognized the homologous morphology between grapevine leaves and leveraged it for purposes of classification and identification. After the European phylloxera crisis, Louis Ravaz proposed using angles of the petiolar veins to classify new resistant rootstocks and the North American *Vitis* spp. from which they are derived (Ravaz, 1902). Pierre Galet extended the measurement of vein angles and ratios of lobe lengths to wine and table grapes, formalizing a system to identify grapevine varieties (Galet, 1979; 1985; 1988; 1990; 2000). Homologous points found in every grapevine leaf lend themselves to geometric and statistical frameworks, allowing the calculation of mean shapes (Martinez and Grenan, 1999) and quantitative genetic studies (Chitwood et al., 2014; Demmings et al., 2019).

**Figure 1:**
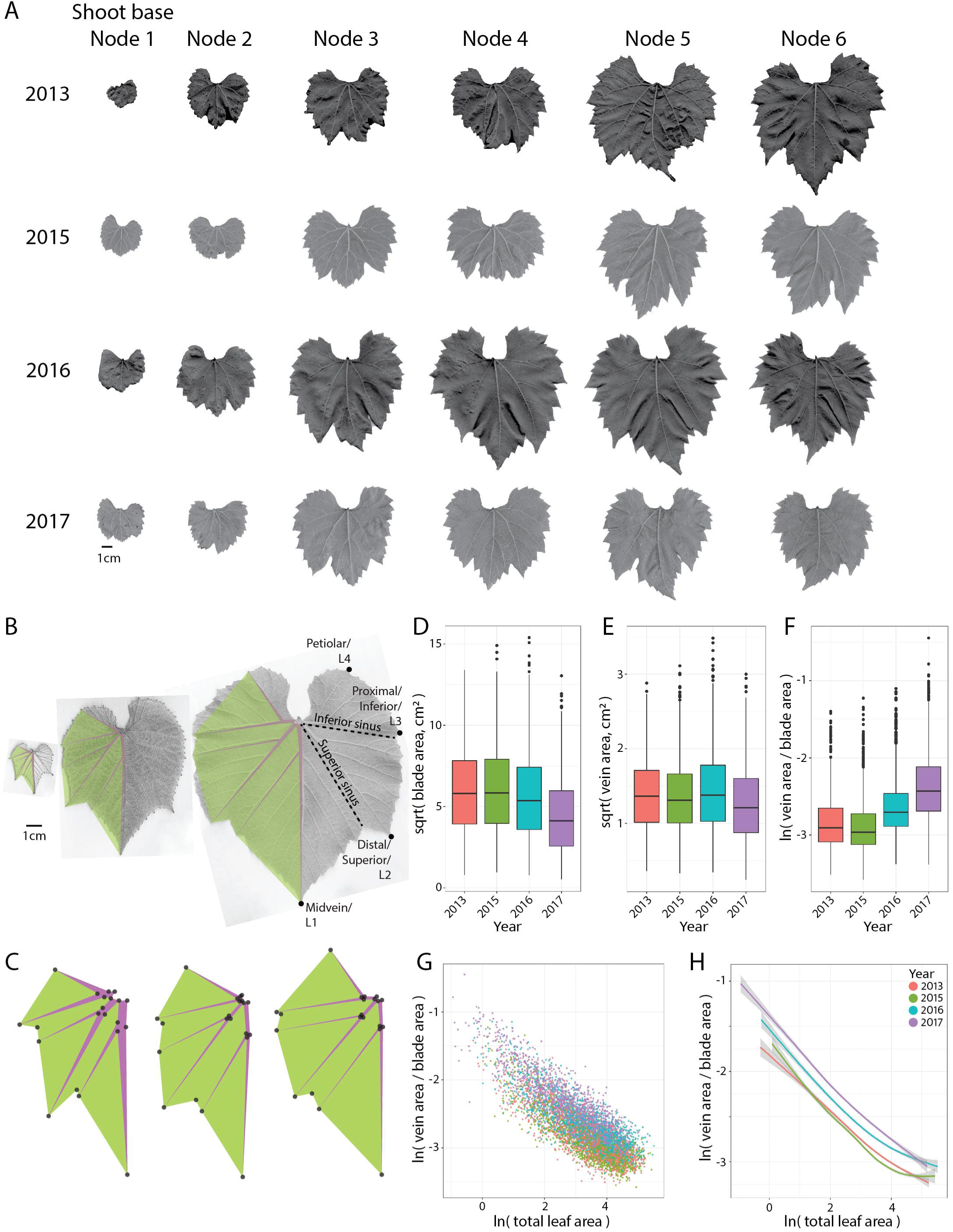
Vein-to-blade ratio is an allometric indicator of leaf size. **A)** For each year, there is a single leaf from each node for each vine sampled. Example leaves matched by node (counting from the shoot base, as these leaves are the first to emerge and most mature) from a *Vitis acerifolia* vine across the four years studied are shown. 1 cm scale is provided. **B)** Leaves from nodes 2, 4, and 6 (left to right) counting from the growing tip of a *V. cinerea* shoot to demonstrate exponential decreases in vein-to-blade ratio as leaves expand. On the left side of each leaf, vein (purple) and blade (green) areas calculated from homologous landmarks are shown. On the right side of the largest leaf, sinuses and lobes tips are indicated with associated ampelometric nomenclature. 1 cm scale is shown. **C)** Landmarks from the leaves in (A) but scaled to comparable areas. Note that the ratio of vein-to-blade area decreases as leaves expand. **D)** Square root of blade area, **E)** square root of vein area, and **F)** natural logarithm of the ratio of vein-to-blade area vs. year as boxplots. **G)** Scatter plot and **H)** loess regression curves (95% confidence limit) for the ln of vein-to-blade ratio vs. the ln of total leaf area. For all panels: 2013, salmon; 2015, green; 2016, turquoise; 2017, purple.

Previously we measured >3,200 leaves sampled from >270 vines of the United States Department of Agriculture (USDA) germplasm repository in Geneva, New York (USA) in 2013 (Chitwood et al., 2016a). From each vine, a complete shoot was sampled, allowing us to record both the node that a leaf arose from and the total leaf count of the shoot. 17 landmarks were used to mark the sinuses, lobes, and vein branch points. In addition to strong species effects, the effects of development were found to independently influence leaf shape. In 2015, we repeated the study, collecting leaves from an entire shoot of the same vines (Chitwood et al., 2016b). This time we increased the number of landmarks to 21 to measure vein width, noting from our first study that the proportional area of the leaf occupied by veins diminishes exponentially as a leaf expands. The distal sinus of leaves from 2015 (the cooler of the two years) was more pronounced than those from 2013, consistent with well-known correlations between serration depth and climate (Bailey and Sinnott, 1915, 1916; Wolfe, 1978; Wilf, 1997; Peppe et al., 2011).

## MATERIALS AND METHODS

### Germplasm, sample collection, and scanning

Leaves were sampled each year from the USDA germplasm repository in Geneva, NY during the second week of June. Eleven species (*Ampelopsis glandulosa* var. *brevipedunculata, Vitis acerifolia, V. aestivalis, V. amurensis, V. cinerea, V. coignetiae, V. labrusca, V. palmata, V. riparia, V. rupestris, V. vulpina*), four hybrids (*V. ×andersonii, V. ×champinii, V. ×doaniana, V. ×novae-angliae*), and 13 vines of *Vitis* spp. with unassigned identity are analyzed in this study. Leaves were collected as stacks in the vineyard keeping track of shoot order and stored in labelled plastic bags with cut holes for ventilation. All leaves greater than ∼1 cm in length starting from the tip of a single shoot were sampled. Leaves were kept in a cooler while collecting in the vineyard and scanned within one to two days. Leaves were arranged on a large format Epson Workforce DS-50000 scanner (Tokyo, Japan) in the order collected from the shoot with a small number near each leaf indicating the node it came from and a ruler for scale. Files are named by the vine ID followed by a sequential lowercase letter if multiple scans are needed to accommodate the leaves of a shoot. We returned to the same vines sampled in 2013 (Chitwood et al., 2016a) and 2015 (Chitwood et al., 2016b) to again sample all leaves from a shoot in 2016 and 2017 (Fig. 1A). Leaves were analyzed using the 21 landmarks that measure vein width in order to measure allometric relationships between vein and blade area (Fig. 1B-C).

### Landmarking

21 landmarks were placed by hand using the point tool as in Fig. 1C on either side of the leaf in ImageJ (Abràmoff et al., 2004). For the 8,412 leaves in this study, 176,652 landmarks were analyzed, or 353,304 values. Landmarks were placed sequentially for each leaf in a scan and saved as a text file of *x* and *y* coordinate values. Landmarks for each scan were visualized using ggplot2 (Wickham, 2016) in R (R Core Team, 2019) to detect mistakes, and if found, the landmarking was redone.

### Data analysis

R scripts (R Core Team, 2019) used for analyzing the data in this study can be found at github: https://github.com/DanChitwood/grapevine_climate_allometry. Spearman’s rank correlation coefficient was calculated using the cor.test() function. All visualizations were made using ggplot2 (Wickham, 2016). In order to correlate leaf shape and size with climate variables without statistical artifacts, leaves were matched such that for each year there is exactly one leaf for each node (counting from the shoot base) from each vine (Fig. 1A). The result is that there is an equal number of leaves represented for each year and that each leaf has a counterpart from the same vine and node in each other year. After matching, the number of leaves analyzed drops from 8,412 leaves to 6,284 and the number of vines analyzed remains 208. Leaf, blade, and vein areas were calculated using landmarks as vertices of a ploygon using the shoelace algorithm, also known as Gauss’s area formula.

Climate data were retrieved from the Northeast Weather Association website via Cornell University (newa.cornell.edu, retrieved September 28, 2019). Daily summaries of minimum, average, and maximum temperatures, leaf wetness hours, and precipitation were analyzed for the Geneva, NY station at the Vineyard North Farm approximately three miles from the USDA germplasm location. For each of the growing seasons, daily data was averaged starting March the previous year (when temperatures begin to rise above freezing in the New York Finger Lakes region) until the beginning of June the following year, approximately one week prior to sampling. Correlation was performed using averages of daily values for the growing season or as averages of a 30 day rolling window, for each calendar day matched across years.

### X-ray Computed Tomography

X-ray CT reconstruction of a *V. riparia* bud was performed using the X3000 North Star Imaging (Rogers, Minnesota, USA) platform and reconstruction software at the Department of Horticulture at Michigan State University.

## RESULTS

For the dataset matched by vine and node (Fig. 1A), blade area varied by year, showing almost identical distributions between 2013 and 2015 and decreasing in 2016 and 2017 (Fig. 1D). Vein area varies little by year (Fig. 1E), consistent with our previous work demonstrating that increases in leaf size are mostly driven by blade expansion. A consequence of this is that the proportion of vein area decreases exponentially relative to blade as a leaf expands (Chitwood et al., 2016b). We plotted the natural log of the ratio of vein-to-blade area by year and found that it varies dramatically relative to blade or vein area alone (Fig. 1F). Vein-to-blade ratio is inversely correlated with leaf size. The relationship is strongly linear when both vein-to-blade ratio and leaf area are natural log transformed (Fig. 1G). The linear relationship between vein-to-blade ratio and leaf area is structured by year, such that each year has nearly identical slopes, indicating allometric relationships between vein and blade area across years are the same (Fig. 1H). However, 2016 and 2017 are shifted to higher vein-to-blade ratio values, reflecting that leaves from these years are smaller than those from 2013 and 2015 (Fig. 1D) and that the differences in leaf size between years are enhanced by vein-to-blade ratio (Fig. 1H).

To determine if there was a relationship between the changes in leaf size and vein-to-blade ratio with climate, we correlated these traits with minimum, average, and maximum daily temperatures and total precipitation and leaf wetness hours as well. Total leaf area, vein area, and blade area weakly correlate with all variables compared to vein-to-blade ratio (Table I). Vein-to-blade ratio is highly positively correlated with maximum daily temperature (rho = 0.482) and negatively correlated with total precipitation and leaf wetness hours (rho = −0.321). The correlations between vein-to-blade ratio with maximum daily temperature and precipitation are consistent across the ten most highly sampled *Vitis* species in the collection (Appendix S1) as well as across nodes (Appendix S2).

**Table I:** Spearman’s rho between leaf traits and averaged climate variables by year.

| Climate variable | Total area | Vein area | Blade area | Vein-to-blade |
| --- | --- | --- | --- | --- |
| Avg. temperature | -0.054 | -0.016 | -0.056 | 0.117 |
| Max. temperature | -0.210 | -0.060 | -0.217 | 0.482 |
| Min. temperature | -0.054 | -0.016 | -0.056 | 0.117 |
| Total precipitation | 0.109 | 0.004 | 0.114 | -0.321 |
| Leaf wetness hrs. | 0.109 | 0.004 | 0.114 | -0.321 |

Median vein-to-blade ratio values for each year increase in the same order as their respective maximum temperature values (Fig. 2A), showing that smaller leaves are associated with high maximum temperature. The effects of temperature on size and shape scale beyond leaves, to shoots, vines, and the vineyard. Looking beyond the dataset matching leaves by node, we considered all leaves from the 208 vines we sampled to calculate leaf count per shoot, total leaf area per shoot, and vineyard leaf area (the total area of leaves sampled). The total leaves per shoot is negatively correlated with maximum temperature (Fig. 2B). The total leaf area of shoots is also negatively correlated with maximum temperature (Fig. 2C), as is expected considering that there are fewer leaves per a shoot with smaller areas at high temperatures. If the total area of leaves sampled for 208 vines is considered, dramatic reductions in overall vegetative growth at the vineyard scale are observed across years. The total leaf area sampled drops from 8.3 m^2^ in 2015 (the coolest year) to 4.4 m^2^ in 2017 (the hottest year), by almost half (Fig. 2D). The same correlations are observed for precipitation, but negative and less strong (Fig. 2E-H), such that smaller leaves are associated with drought. Although vein-to-blade ratio is a subtle morphological feature of leaves that amplifies differences in leaf area (Fig. 1D, 1H), such differences have large effects when scaled to the levels of vine and vineyard (Fig. 2).

**Figure 2:**
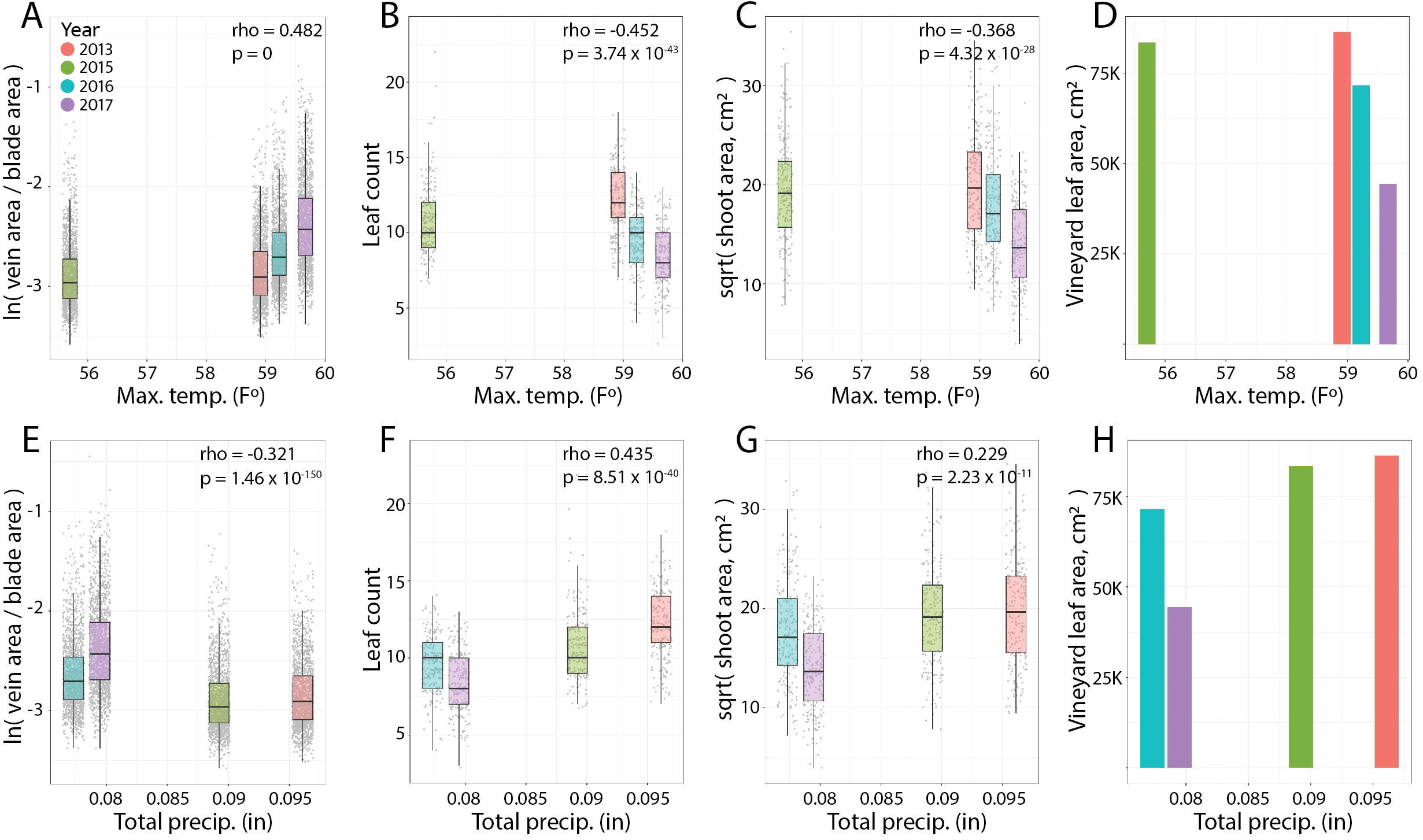
Correlation of leaf shape, size, and count and total shoot and vineyard leaf areas with maximum temperature and precipitation. **A)** ln of vein-to-blade ratio, **B)** leaf count per shoot, **C)** square root of total leaf area per shoot, and **D)** total leaf area for all shoots sampled (vineyard area) vs. average maximum temperature for each year as boxplots. **E-H)** The same as above but plotted against average precipitation for each year. For all panels: 2013, salmon; 2015, green; 2016, turquoise; 2017, purple.

The above correlations average climate variables over a 15 month period, from the previous March when vines begin to thaw to June the following year when leaves were harvested. This is because 5-7 leaf primordia are pre-formed in buds, initiating the previous year from when they emerge. The multiyear development of grapevine primordia is better described for inflorescences (Srinivasan and Mullins, 1976), which are conspicuously present in grapevine buds (Fig. 3A). The number of pre-patterned inflorescences is influenced by the environment of the previous year (Khanduia and Balasubrahmanyam, 1972; Guilpart et al., 2014) and can even be used to predict cluster counts for the coming year (Vasquez and Fidelibus, 2006). As we matched nodes by mature leaves counting from the base of the shoot, a large proportion of the leaves analyzed in this study were not neoformed the year of harvest and potentially influenced by the climate of the previous year in which they initiated.

**Figure 3:**
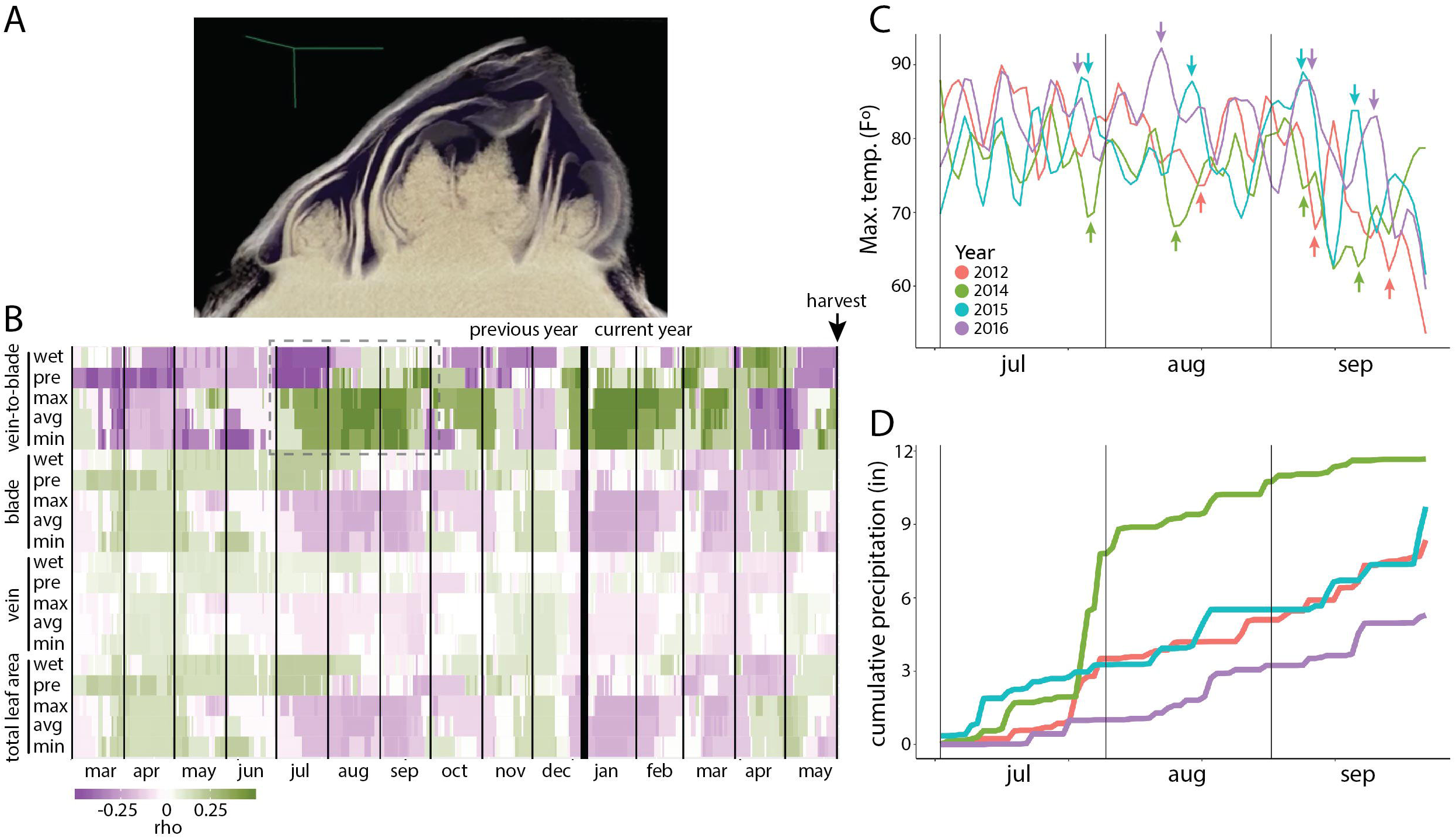
Correlations between leaf size and shape with maximum temperature and precipitation during leaf development. **A)** X-ray Computed Tomography (X-ray CT) 3D reconstruction of a bud from *Vitis riparia*. Note cluster and leaf primordia of primary and secondary meristems that had formed the previous growing season and will grow and expand the following year. **B)** Correlation of total leaf area, vein area, blade area, and vein-to-blade ratio with daily values of leaf wetness hours (wet), precipitation (pre), maximum temperature (max), average temperature (avg), minimum temperature (min) across four years. Spearman’s rho values (purple to green, negative to positive) are shown as a 30 day sliding window for each day starting March the previous year of harvest to the harvest time of the following year at the beginning of June. Dotted box indicates a period of high correlation the summer of the previous year. **C)** Plot of maximum daily temperatures for the previous summer months. Extreme temperature events are indicated by colored arrows. **D)** Cumulative precipitation for previous summer months. Curves are colored by year. For all panels: 2012-2013, salmon; 2014-2015, green; 2015-2016, turquoise; 2016-2017, purple.

To pinpoint which period of time during leaf development climate exerts the strongest influence on morphology, we correlated daily climate values as a sliding window with total leaf area, vein area, blade area, and vein-to-blade ratio (Fig. 3B). Again, vein-to-blade ratio is considerably more correlated with climate variables than leaf, blade, or vein area alone. Considering that during the winter buds are frozen (Londo and Kovaleski, 2017; Kovaleski et al., 2018; Londo and Kovaleski, 2019; Kovaleski and Londo, 2019), the most conspicuous window of vein-to-blade ratio correlation in the correct direction (positive correlation with temperature and negative correlation with precipitation) is the previous summer. This does not discount environmental influences from bud burst in spring of the current year to harvest, but previous year climate trends are consistent with a role in influencing leaf size and shape. For example, the two coolest growing seasons (2012-2013 and 2014-2015) and the two warmest growing seasons (2015-2016 and 2016-2017) have extreme cold and heat maximum temperature events, respectively, the previous summer (Fig. 3C). Similarly, the coolest growing season (2014-2015) is also by far the wettest previous summer, and the hottest growing season (2016-2017) was the driest previous summer (Fig. 3D). In fact, 2016 was a historic drought in the Finger Lakes region, centered in Geneva, New York (USA) where the sampled USDA germplasm repository is located (Sweet et al., 2017).

## DISCUSSION

We find that the ratio of vein-to-blade area is an allometric indicator of leaf size that varies more strongly by year than vein or blade area alone (Fig. 1). Vein-to-blade area correlates most strongly with maximum daily temperature and precipitation, such that smaller leaves are associated with heat and drought (Table I). The effects of climate on leaf morphology propagate to larger scales with implications for grapevine vigor and their cultivation. Total leaf count per shoot, total shoot leaf area, and total leaf area studied (vineyard area) all negatively correlate with maximum temperature and positively correlate with precipitation (Fig. 2). These correlations are strongest for the summer previous the year of harvest when grapevine primordia initiate in buds before winter dormancy and are consistent with extreme climate events at the study location (Fig. 3).

That climate would alter grapevine morphology is not unanticipated. The environmental sensitivity of grapevines to climate variability is long recognized and has been measured in many ways. Several unique developmental, life history, and cultivation characteristics of grapevines lend themselves to the study of the relationship between plants and climate (Carmona et al., 2008). Records of grapevine harvest dates going back centuries exist because of the cultural importance of wine to Europeans and have been correlated with (and used to predict) past temperature anomalies (Chuine et al., 2004; Menzel et al., 2005; Meier et al., 2007). This is possible because many of the same wine grape varieties that existed centuries ago still exist today because of clonal propagation, permitting historical longitudinal analyses. Similarly, predictions of the impact of future climates on grapevine growth and wine quality have been made (Jones et al., 2005; De Orduna, 2010; Hannah et al., 2013). Because grapevines are long-lived perennials, typically harvested over longer periods than other crops, larger investments are made when choosing the location to establish a vineyard.

The long-term economic implications of climate change for grapevines and the wine industry are evident in the very data presented in this study. We can report effects of precipitation because the vineyard we sampled is not irrigated. An analysis of the effects of the historic 2016 New York drought reveals that rainfed crops in the region needed to increase their water use by 500-600% (Sweet et al., 2017). From this perspective, unlike annuals, grapevines remain a constant as the climate changes around them. As the grapevine phenotype is a summation of past and current environmental effects, it can be leveraged to study how plants respond to climate change. However, this is also a liability to viticulture and the wine industry. Even over the short-term, the effects of climate linger longer than in annuals, and the environment of the previous year cannot be discounted when studying annual variation, yield, or crop quality in grapevines (Khanduia and Balasubrahmanyam, 1972; Vasquez and Fidelibus, 2006; Guilpart et al., 2014).

The ability to calculate vein-to-blade ratio is permitted by the unique morphology of grapevine leaves that contain a multitude of homologous landmarks found across nearly all *Vitis* species (Fig. 1B-C). The use of landmarks in grapevine leaves is inspired by ampelometry, a more than century old tradition of using geometric morphometrics to quantify and classify grapevine varieties (Ravaz, 1902; Galet, 1979; 1985; 1988; 1990; 2000). Yet, despite the uniqueness of the approaches presented here to grapevines, our results have implications for the relationship between plants and climate in general. The correlation between leaf size and toothed margins with the climate has long been studied in present day and ancient plants (Bailey and Sinnott, 1915, 1916; Wolfe, 1978; Wilf, 1997; Peppe et al., 2011). For both these traits, allometry plays an important role. Leaf size is of course fundamental to allometric relationships, and a theory of its functional significance is well established (Parkhurst and Loucks, 1972). Many hypotheses have been put forward regarding the functional significance of teeth (Nicotra et al., 2011), but as vascularized extensions of the margin that are predominate in leaf primordia (Jones et al., 2013) and diminish in relative size as the leaf expands, allometry and specifically vein-to-blade ratio are predominate morphometric features. Lobes and teeth might be a developmental constraint resulting from the packing of leaves in buds, as is the case for grapevine (Edwards et al., 2016). Recently, the allometry of teeth for the grapevine leaves from the 2013 and 2015 growing season described in this study was undertaken, noting their phenotypic plasticity across years, developmental context, and the meaningfulness of measuring traits used for paleoclimate reconstructions in a living germplasm collection (Baumgartner et al., 2020). The relationship between vein and blade is a fundamental feature of all leaves, an indicator of development itself that is modulated by both evolutionary and environmental forces.

## CONCLUSIONS

By tracing the movement of pin pricks arranged as a grid on a young fig leaf, Stephen Hales documented the allometric growth of leaves in 1727: “By observing the difference of the progressive and lateral motions of these points in different leaves, that were of very different lengths in proportion to their breadths” (Hales, 1727). Here, we document a strongly linear allometric relationship between the ratio of vein-to-blade area with grapevine leaf size. Vein-to-blade ratio in grapevine leaves is more strongly correlated with climate variables than leaf size itself. Blade area expands exponentially faster than vein area, making it a sensitive indicator of leaf size and shape. Using vein-to-blade ratio as an allometric indicator, we document the effects of climate on grapevine leaf morphology across 208 vines of North American *Vitis* species in a four year study, showing that drought and heat of the previous year reduce leaf size. Our results show how climate directly alters the morphology of grapevines through allometry and has implications for monitoring the relationship between plants and climate in the past, present, and future.

## ACKNOWLEDGEMENTS

DHC and JPL were recipients of a Grape Research Coordination Network stipend from the National Science Foundation that funded this work. DHC, ZM, and JPL were supported by the National Science Foundation Plant Genome Research Program award number 1546869. This project was supported by the USDA National Institute of Food and Agriculture, and by Michigan State University AgBioResearch.

## AUTHOR CONTRIBUTIONS

DHC and JPL conceived of the project and experimental design. JM, ZM, and MF collected data. DHC and RV designed statistical analyses. DHC analyzed data and wrote the paper with input and editing from JM, ZM, MF, RV, and JPL.

## DATA AVAILABILITY

All data and code used in this study can be found on github (https://github.com/DanChitwood/grapevine_climate_allometry).

## SUPPORTING INFORMATION

Appendix S1: Spearman’s rho between leaf traits and averaged climate variables by year for each species.

Appendix S2: Spearman’s rho between leaf traits and averaged climate variables by year for each node from the shoot base.

